# Evolutionary emergence of alternative stable states in shallow lakes

**DOI:** 10.1101/2022.02.23.481597

**Authors:** Alice N. Ardichvili, Nicolas Loeuille, Vasilis Dakos

**Affiliations:** Sorbonne Université, Université de Paris-Cité, UPEC, CNRS, INRA, IRD, UMR 7618, Institute of Ecology and Environmental Sciences – Paris, France; Université de Montpellier, IRD, EPHE, CNRS, UMR 5554, Institut des Sciences de l’Evolution de Montpellier – Montpellier, France

**Keywords:** tipping points, bistability, adaptive dynamics, macrophyte, phytoplankton, diversification, eco-evolutionary model

## Abstract

Ecosystems under stress may respond abruptly and irreversibly through tipping points. Although much is explored on the mechanisms that affect tipping points and alternative stable states, little is known on how ecosystems with alternative stable states could have emerged in the first place. Here, we explore whether evolution by natural selection can lead to the emergence of ecosystems with tipping points using a toy model inspired by shallow lakes. Shallow lakes are the best-known example where a shift from a submersed macrophyte dominated state to a floating macrophyte dominated state in response to excess nutrient loading corresponds to a tipping point between alternative stable states. We model the evolution of the macrophyte’s living depth in the lake, we identify the conditions under which the ancestor macrophyte population diversifies, and investigate whether alternate stable states dominated by different macrophyte phenotypes occur. Given the trade-off between access to light and nutrient along the water column, we show that asymmetry in competition for light is required for diversification, while alternative stable states require an additional mechanism, an unequal efficiency in nutrient removal. We find that eco-evolutionary dynamics may produce alternative stable states, but only under a restrictive range of conditions. Our analysis suggests that competitive asymmetries along opposing resource gradients may allow alternative stable states to emerge by natural selection.

## Introduction

Ecosystems are exposed to increasing stress of habitat fragmentation, biotic exploitation, climate change, and pollution (Intergovernmental Science-Policy Platform on Biodiversity and Ecosystem Services, 2019). Under such stress, ecosystems could possibly respond in an abrupt, unpredictable, and irreversible way -that is a tipping point-, which may be a concern for future conservation (Scheffer, Carpenter, et al., 2001). Theoretically, evolution may affect the stability of ecosystems (Kondoh, 2003; Loeuille, 2010). In particular, evolution by natural selection could prevent extinction of a species, a process called evolutionary rescue (Gomulkiewicz and Holt, 1995). Dakos et al., 2019 have postulated that trait variation could alter the bistability of an ecosystem, either by hastening the approach to a tipping point, delaying it, or even removing bistability. However, could it be that natural selection, instead of acting like a rescuer, allows the emergence of the very dynamics that threaten ecosystems with tipping points?

In ecosystems with tipping points, strong positive feedbacks are responsible for the establishment of alternative stable states (Nes, Arani, et al., 2016). Alternative stable states correspond to at least two equilibria that are locally stable in a given range of environmental conditions. The transition from one stable equilibrium to another is abrupt: a small environmental perturbation past a threshold can push the ecosystem into a qualitatively different state (Holling, 1973). The ecosystem also exhibits hysteresis: once the ecosystem has shifted to an alternative stable state, restoring environmental conditions to the threshold when the collapse occurred is not sufficient to return the system to its previous equilibrium state (Scheffer, Carpenter, et al., 2001). Hysteresis makes the management of ecosystems with alternative stable states difficult since a large restoration is needed to return to the pre-collapsed (and often desirable) state. Examples of ecosystems with alternative stable states include drylands which can shift from shrub vegetation to desert (Kéfi et al., 2007), tropical forests that can switch from forest to savanna and back (Scheffer, Vergnon, et al., 2014; Staver et al., 2011), and coral reefs that can shift from a healthy coral dominance to being overgrown by macro-algae (Knowlton, 1992). Yet, the most studied examples of tipping points between alternative states are found in shallow lakes (Gsell et al., 2016).

In shallow lakes, tipping responses can be found between two alternative stable states dominated by either submersed or floating macrophytes, that is two different types of vascular aquatic vegetation. The positive feedback responsible for alternative stable states is created by three mechanisms (Scheffer, Szabo, et al., 2003): 1) a light-nutrient trade-off that makes submersed macrophytes better competitors for nutrients and floating macrophytes better competitors for light, 2) an asymmetric competition for light caused by the shading of floating macrophytes, 3) a difference in the efficiency at removing nutrients from the water column. These three mechanisms allow better access of submersed macrophytes to nutrients than floating macrophytes, so that submersed macrophytes remove nutrients from the water and maintain a clear water state. When nutrient loading exceeds a certain threshold, floating macrophytes are able to grow from surplus nutrients that can no longer be retained by submersed macrophytes. In addition, the ability of floating macrophytes to shade submersed macrophytes hinders submersed macrophytes growth. As a result, more nutrients become available to floating macrophytes, floating macrophytes grow more, further shade, and eventually outcompete the submersed macrophytes, and the water clarity in the lake decreases. Because of this positive feedback, a large reduction in nutrient loading is required to restore the dominance of submersed macrophytes and the clear water state in the lake.

Evolution can interact with the stability of ecosystems in at least three ways: by changing their persistence (number of species lost), resistance (amount of environmental change that a system can take before shifting to another state), and resilience (return time to equilibrium). Models of evolutionary rescue suggest that a population’s heritable variability may promote its persistence in a changed environment (Gomulkiewicz and Holt, 1995). Theoretical work also showed that a forager’s evolution enables the persistence of species in complex food webs (Kondoh, 2003). In larger communities, Cortez et al., 2020 found contrasting effects of eco-evolutionary feedbacks: evolution of exploiter species’ traits tends to prevent loss of species, while evolution of exploited species tends to do the opposite. In that sense, evolution increases persistence at the ecosystem level. A theoretical study showed that evolution tends to increase the resilience of small communities by shortening their return time to equilibrium, while evolution tends to decrease the resilience of large communities (Loeuille, 2010). However, only recently are we beginning to examine whether evolution could also alter the stability properties of ecosystems with tipping points (Chaparro Pedraza et al., 2021; Dakos et al., 2019).

It is reasonable to hypothesise that evolution could affect the presence of dynamic attractors responsible for alternative stable states, for instance affecting the asymmetry of competition processes or the strength of positive interactions that are at the very basis of some positive feedback loops. Chaparro-Pedraza and Roos, 2020 showed that a minor ecological change can trigger a future tipping point. In that study, a momentary decrease in mortality triggered the evolution of body size, which resulted in a population shift in the long run (Chaparro-Pedraza and Roos, 2020). This study focuses on the impact of evolution in ecosystems with tipping points, but the role of evolution in the emergence of positive feedbacks that are responsible for the existence of tipping points is, to our knowledge, unaddressed.

So how could ecosystems with tipping points have emerged in the first place? Here, we hypothesize that the evolution of a population’s trait could explain the emergence of strong positive feedbacks that eventually lead to tipping points. The literature that looks at ecosystems with tipping points from an ecological perspective, without evolution, examines the conditions that are necessary for alternative stables states to occur in different systems (De Roos and Persson, 2002; Kéfi et al., 2007; Mumby et al., 2007; Scheffer, Szabo, et al., 2003). However, we do not know if and under which conditions alternative stable states will emerge in an ecosystem after its long term evolution. Put differently, the question is: when does evolution lead to the emergence of the strong positive feedbacks needed for the observed tipping points in ecosystems around us?

In this paper, we tackle this question in the particular case of alternative states between submersed and floating macrophytes in a shallow lake. We model the adaptive dynamics of a macrophyte population whose growth depth in a shallow lake can evolve. A macrophyte’s growth depth in the lake can be under the genetic control of the length of its stem or the buoyancy of its leaves. We first study the conditions under which a macrophyte population can diversify. Diversification is required for alternative states to emerge because at least two phenotypes are needed to dominate alternatively. We expect that the way nutrients and light are distributed in the water column and the type of competition between diversified phenotypes should be a strong determinant of evolutionary outcomes. Second, we study how the evolved community responds to increased nutrient loading. If coexisting phenotypic populations are indeed differentiated in terms of competitive ability for light and nutrients, this competitive ability should determine the response of the lake ecosystem to perturbations, eventually exhibiting tipping points. We hypothesize that in sufficiently differentiated phenotypic populations, asymmetries in competitive abilities for light and nutrients should create strong positive feedbacks that could lead to the establishment of alternative stable states.

We explore these questions using three different evolutionary scenarios combining three mechanisms present in a shallow lake (Fig. 1). In the first scenario, depth determines the quantity of light and nutrients that a macrophyte population has access to. In the second scenario, we add asymmetric competition created by shading: shallower macrophyte populations shade deeper populations. Finally, in the third scenario, we add model a higher nutrient exploitation efficiency for macrophyte populations that grow closer to the sediment. In what follows, we first define the ecological dynamics of a single macrophyte population, we describe the ecological mechanisms that can affect evolutionary trajectories in the macrophyte populations, and we then present results structured along the three described evolutionary scenarios.

**Figure 1.**
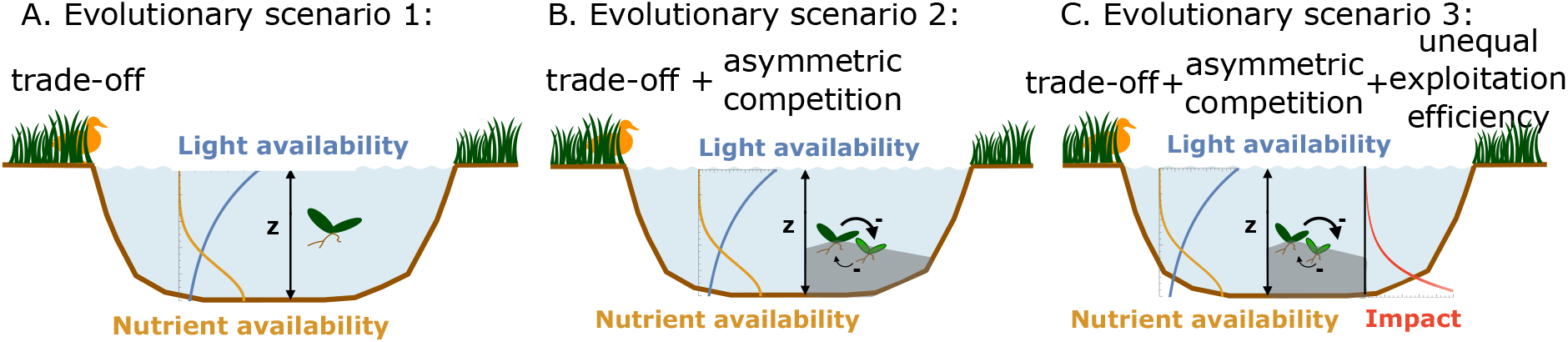
Three evolutionary scenarios based on different combinations of ecological mechanisms in a shallow lake model. A: in scenario 1, the depth at which a macrophyte population settles is simply associated with a trade-off between access to light and nutrients. B: in scenario 2, in addition to the trade-off, we introduce asymmetric competition for light, i.e. a phenotype that grows higher in the water column shades all phenotypes below it. C: in Scenario 3, on top of the two previous mechanisms, we consider different nutrient exploitation efficiencies whereby phenotypes that grow deeper in the water column are more efficient at removing nutrients. (z denotes water column depth, arrows the light competition between macrophyte phenotypes, impact is the effect of macrophytes on nutrient content in the water column)

## Material and methods

We model the eco-evolutionary dynamics of a photosynthetic organism whose depth in the water column of a lake, *z*, evolves. As exemplified by Scheffer, Szabo, et al., 2003, alternative stable states can exist between floating vs submersed macrophytes (Scheffer, Szabo, et al., 2003), between green algae and cyanobacteria dominance (Bengfort and Malchow, 2016), as well as between submersed macrophytes and phytoplankton (Scheffer, 1998). The depth at which a rooted macrophyte lives in the lake can be thought of as the height that its canopy reaches, which is controlled by the length of its stem. Depending on their size, buoyancy and propensity to form colonies, different phytoplankton species can also settle at different depths (Alexander and Imberger, 2008). Thus, in these systems, depth is an important trait that determines the competitive ability and potential asymmetries between phenotypes.

Depth in the lake, *z*, varies from the surface (*z* = 0) to the bottom (*z* = *z*_*b*_). In the rest of the paper, small values of *z* are closer to the surface and are referred to as “shallower” or “above”. Larger values of *z* are deeper and referred to as “below”.

### Ecological dynamical model of a macrophyte population

We modeled the growth rate of a single macrophyte population following Scheffer, Szabo, et al., 2003:

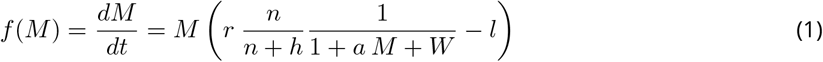

with

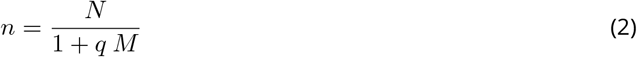

Change in a macrophyte population biomass, *M*, over time depends on maximum intrinsic growth rate *r*, which is limited by nutrient and light availability, and on mortality rate *l*. Light limitation and nutrient limitation are two mutliplied Monod terms. Light limitation depends on *W*, the background turbidity of the water, and the shading induced by conspecifics, *aM*. Nutrient limitation depends on the amount of available nutrients in the water column, *n*, with half-saturation constant *h*, which represents the macrophyte’s competitive ability to acquire nutrients. A low *h* implies that macrophytes are able to grow at low nutrient levels, while a high *h* means that nutrients quickly become limiting; a larger amount of nutrient is required to sustain macrophyte growth. We do not model the nutrient dynamics, but the amount of nutrients available in the water *n* depends on the quantity of nutrients *N* that would be present in the absence of macrophytes, and on the density of macrophytes that uptake nutrients from the water at rate *q*. A list of parameters is presented in Table 1.

**Table 1.** Variables and parameters, their meaning, unit, and default values

|  | Meaning | Unit | Default value |
| --- | --- | --- | --- |
| $t$ | time | day | - |
| $M$ | macrophyte biomass | $\text{g L}^{-1}$ | - |
| $z$ | macrophyte depth in the lake, the evolving strategy | m | - |
| $r$ | maximum macrophyte growth rate | $\text{day}^{-1}$ | 0.5 |
| $l$ | macrophyte mortality rate | $\text{day}^{-1}$ | 0.05 |
| $h$ | macrophyte dependency on water nutrient | $\text{mg L}^{-1}$ | 0.1 |
| $N_0$ | total amount of nutrient released from the sediment | $\text{mg m}^{-2}$ | 5 |
| $u$ | strength of nutrient diffusion | m | - |
| $W_0$ | baseline light attenuation | dimensionless | 1 |
| $w$ | strength of light attenuation | $\text{m}^{-1}$ | - |
| $a_0$ | intra-specific light competition coefficient | $(\text{g/L})^{-1}$ | 0.01 |
| $b$ | strength of asymmetry in light competition | $\text{m}^{-1}$ | 1 |
| $q_0$ | surface macrophyte impact on water nutrient content | $(\text{g/L})^{-1}$ | 0.05 when $p=0$<br>0.005 otherwise |
| $p$ | difference in nutrient exploitation efficiency | $\text{m}^{-1}$ | 0.5 |

**Table 2.**
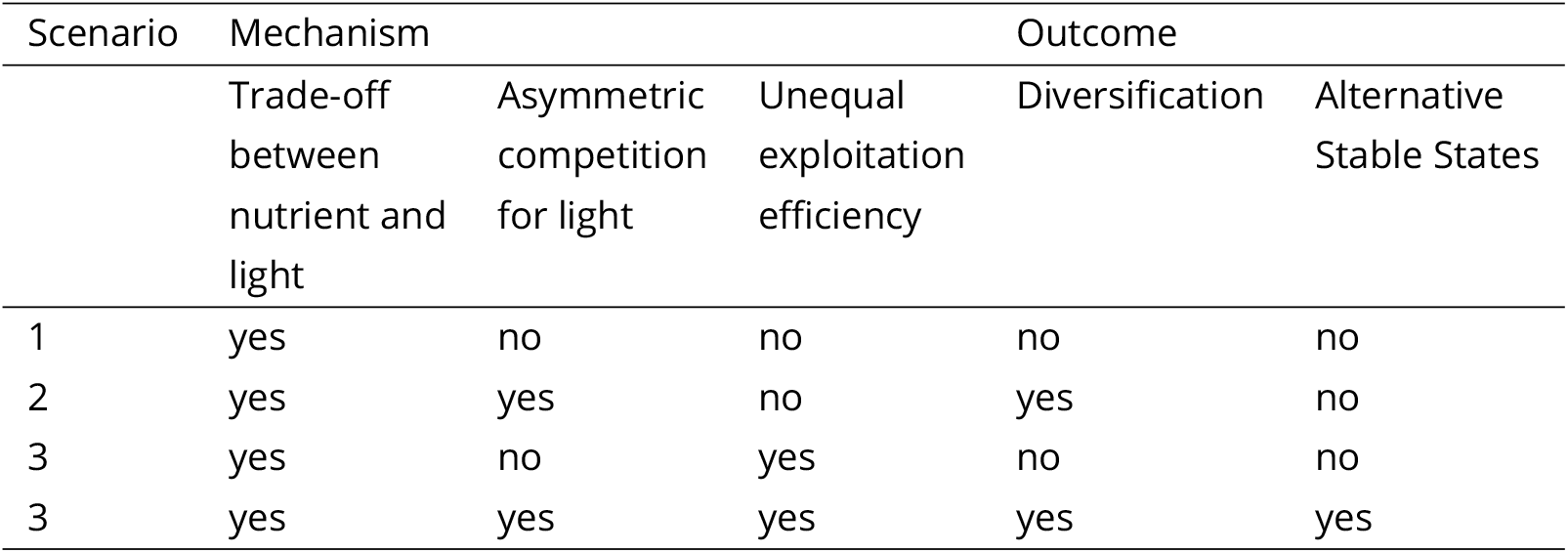
Summary of the mechanisms of each scenario and their result

We detail below the three ecological mechanisms used to explore the three evolutionary scenarios presented in Fig. 1. These scenarios imply that parameters *N, W, a*, and *q* depend on the evolving trait, depth in the lake, *z*.

### Evolutionary analysis using adaptive dynamics

We analyse the evolution of a macrophyte population using adaptive dynamics (AD) (Dieckmann and Law, 1996; Geritz et al., 1998; Metz et al., 1992). In the adaptive dynamics framework, each population is characterized by a quantitative phenotypic trait (in our case *z*, depth in the lake). The main assumption of AD is the separation of evolutionary and ecological timescales: the dominant population with trait *z* (called the resident population) is assumed to be at its ecological equilibrium and sets the conditions felt by invading phenotypes (called mutant populations, characterized by trait *z*_*m*_). The dynamics of the interacting resident and mutant populations are:

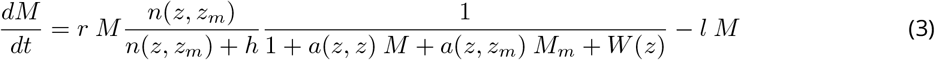

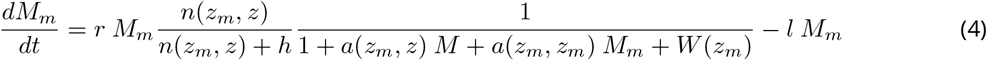

where

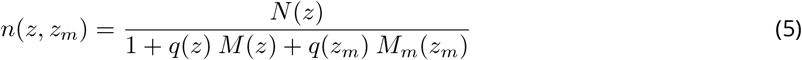

The expressions of *W* (*z*), *N* (*z*), *a*(*z, z*_*m*_), and *q*(*z*) depend on the scenario in which the population evolves. *n*(*z*_*m*_, *z*) is symmetrical to *n*(*z, z*_*m*_).

The introduced mutant is assumed to be rare (*M*_*m*_ *≈* 0) while the resident has time to reach its equilibrium (*M* ^*∗*^). Whether a mutant is able to invade the resident population depends on the sign of the invasion fitness, defined as the *per capita* growth rate of a rare mutant in the stationary conditions set by the resident population of trait value *z*. By definition, identical phenotypes can neither invade nor be invaded by the resident population and have a null invasion fitness: *s*(*z, z*) = 0. Mutants with a negative invasion fitness die out. Mutants with a positive invasion fitness successfully invade the resident population. The invasion fitness *s*(*z*_*m*_, *z*) of the mutant is:

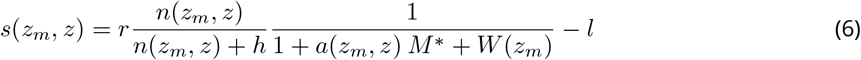

with

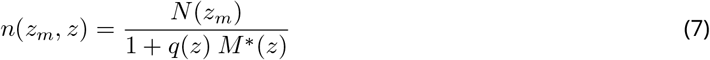

From the invasion fitness equation, we derive the fitness gradient and find long term evolutionary outcomes, or Evolutionary Singularities (ES). ES are classified by deriving the convergence and invasibility criteria from fitness second derivatives (Geritz et al., 1998).

### The three ecological mechanisms explored in the three evolutionary scenarios

We now describe the three ecological mechanisms used to explore the three evolutionary scenarios presented in Fig. 1.

### Vertical trade-off for light and nutrients (evolutionary scenario 1, 2 and 3)

In every scenario, we model light attenuation under water following the Beer-Lambert law as in (Huisman and Weissing, 1995), and assume that nutrients are stored in the sediment and brought in the water column by eddy diffusion as in Klausmeier and Litchman, 2001. These assumptions translate into the expression of *W* and *N* as functions of *z* (Fig. 2A and B) :

**Figure 2.**
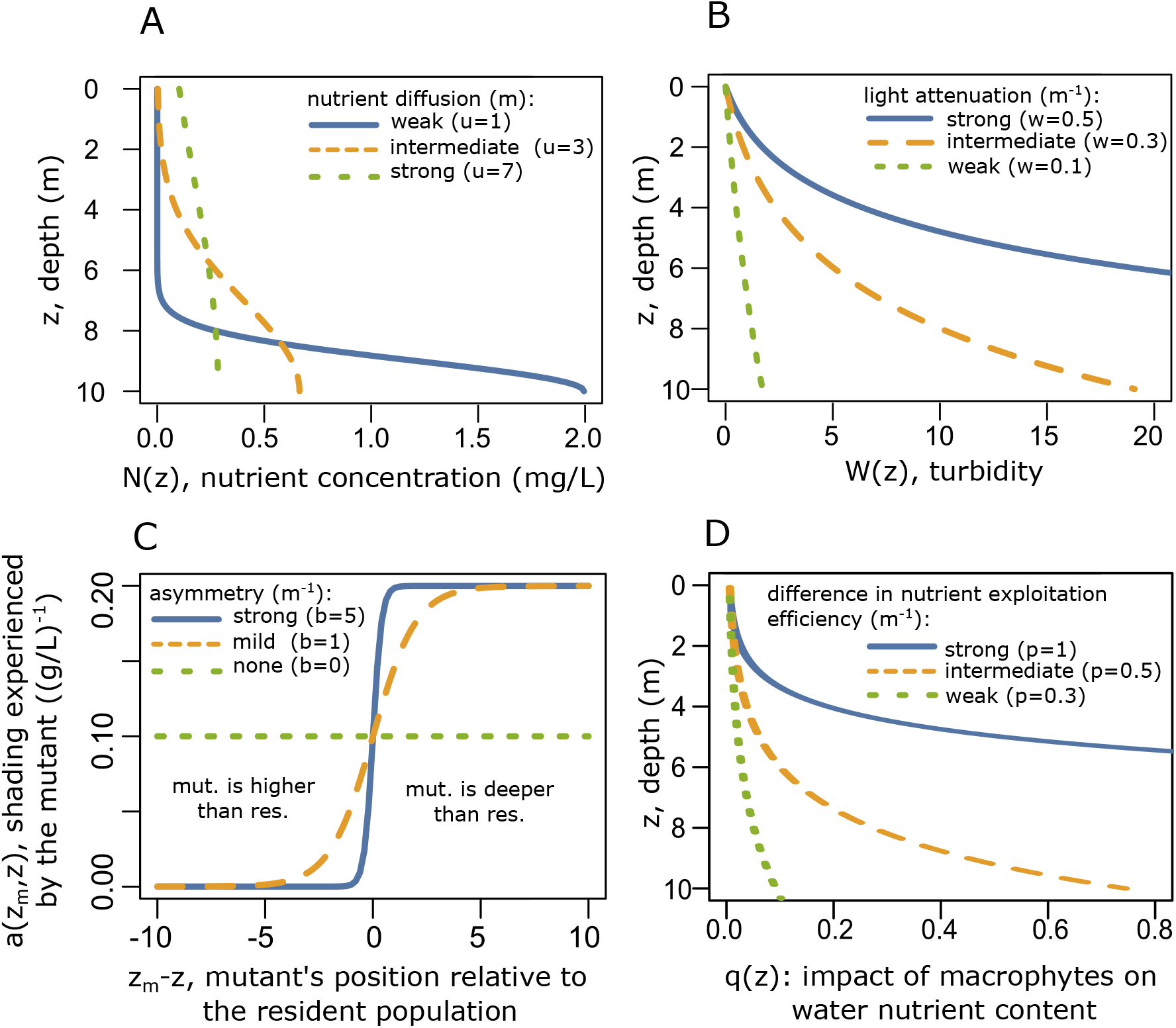
Functions used to model each mechanism. A: Trade-off between nutrients and light: nutrients are diffused from the bottom of the lake with strength *u*, but the total quantity of nutrients in the water is constant (*N*_0_ = 5*mg/m*^2^) B: Trade-off between nutrients and light: turbidity increases with depth at a different pace depending on *w*, the strength of attenuation (*W*_0_ = 1) C: Asymmetric competition by shading: parameter *b* controls how steep the asymmetric competition between two populations is (*a*_0_ = 0.01(*g/L*)^*−*1^) D: Unequal exploitation efficiency: Macrophyte’s ability to retain nutrients increases with depth (*q*_0_ = 0.005(*g/L*)^*−*1^

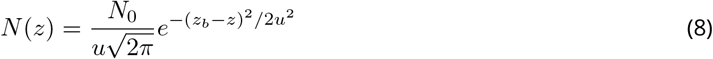

*N*_0_ is the total amount of nutrient in the water and *u* controls the strength of nutrient diffusion (Fig. 2A). A high *u* implies that nutrients are more homogeneously distributed, while a low *u* corresponds to strong stratification of nutrient in the water column. The total quantity of nutrients *N* is constant, regardless of diffusion (*u*) variation.

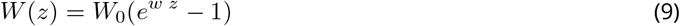

*W*_0_ is the baseline turbidity and *w* controls the strength of light attenuation (Fig. 2B). A low *w* implies that luminosity barely changes with depth *z*, while a high *w* means that the bottom of the lake is much darker than the surface.

### Asymmetric competition by shading (evolutionary scenario 2,3)

In addition to the vertical trade-off for light and nutrients, we consider a second mechanism, that of asymmetric competition for light (ie. shading, Fig. 2C). We model shading by assuming that the shading coefficient *a* varies with the relative depth between two populations :

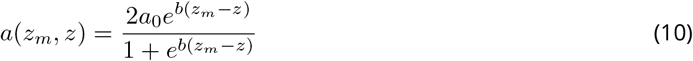

This variation captures the strength with which a population settling deeper is shaded by the shallower populations. In that sense, competition for light is asymmetric as only the above population has an effect on the population below it in the water column. The intensity of such asymmetric competition for light depends on the depth difference between the two competing populations and the sensitivity parameter *b* (Fig. 2C). We study this mechanism in the second and third evolutionary scenarios.

### Unequal nutrient efficiency (evolutionary scenario 3)

The third mechanism introduces the fact that macrophytes living at different depths have different competitive abilities for nutrient. (Fig. 2D and 11). Deeper macrophytes remove nutrients more effectively than shallower macrophytes from the water column, because they prevent the re-suspension of sediment by intercepting nutrients diffusing from the bottom with their shoots. We include mechanism in scenario 3 by letting the parameter *q* increase with macrophyte depth *z*:

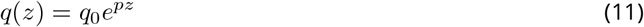

This expression implies that deep macrophytes have a stronger impact on water nutrient content than shallower macrophytes. Setting *p* = 0 is equivalent to turning off this difference in exploitation efficiencies, which is the case in scenario 1 and 2 (Fig. 2 D).

### Identifying conditions for diversification

One type of ES in our model is a convergent and non-invasible strategy, called a Continuously Stable Strategy (CSS). It means that the macrophyte population converges to a selected depth. The other ES we obtain is a Branching Point (BP): that is an ES that is convergent but invasible. It means that the macrophyte population evolves to a certain depth at which disruptive selection acts and polymorphism emerges. Evolution to a BP leads to diversification and thus satisfies the first condition for the emergence of a system with alternative stable states: the existence of at least two phenotypes.

The sign plot of the invasion fitness eq. 6 according to the mutant trait *z*_*m*_ and the resident trait *z* is a Pairwise Invasibility Plot (PIP), and is used to visualise long term ES. Although it is not possible to obtain an analytical expression of the ES *z*^*∗*^, we numerically computed the value of *z*^*∗*^ for different combinations of nutrient diffusion *u* and light attenuation *w* using the invasion fitness equation. We also present how the ES varies with environmental conditions in *E*^3^-diagrams (Ferriere and Legendre, 2013), which are plots showing the ES as a function of parameters *u, w, b* and *p*. The range of parameters used was sufficiently large to capture extreme strategies (macrophyte population evolving to the bottom or to the surface of the lake).

When diversification is possible (BP), no mathematical analysis can be used to predict if further diversification can happen nor how many phenotypes can eventually coexist. Therefore, we present simulations of the eco-evolutionary dynamics for combinations of parameters that yielded a branching point. The parameters for which we ran simulations were sampled at regular intervals of *u* and *w* to cover the diversification region. Simulations are time discrete. They start with a single ancestor phenotype (*z* = 1) at its equilibrium density. At every time step, potential parent populations mutate with a rate of 10^*−*4^ times their density. If the parent population does mutate, the mutant trait is drawn from a normal distribution centered around the parent trait with a standard deviation of 0.05 meters. The mutant is then introduced with an initial density of 10^*−*2^ g/L. The mutant population grows depending on its population dynamics described in eq 3,4, experiencing light and nutrient competition from all other populations. Populations falling below the density of 10^*−*2^ g/L are considered extinct. At the next time step, populations above that threshold are potential candidates for generating mutants, and evolution proceeds with the sequential replacement of populations.

### Identifying the emergence of alternative stable states after evolution

To assess whether evolution leads to the emergence of alternative stable states, we model the impact of an ecological perturbation: nutrient enrichment. We assume that the population trait *z* does not evolve in response to the perturbation, but only its equilibrium density is affected. Specifically, after the eco-evolutionary simulations reach a quasi-equilibrium (when trait *z* values did not change by more than 0.1 meters during 2 *∗* 10^7^ timesteps), we retrieve the value of the depth *z*_1_ and *z*_2_ of the two phenotypes. We then analyse their corresponding ecological dynamics using the same model as described before, with the growth of the phenotypes *i* depending on the phenotype *j*:

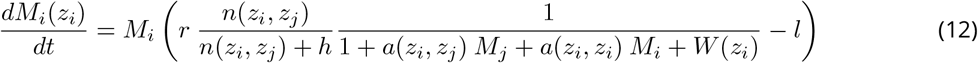

With

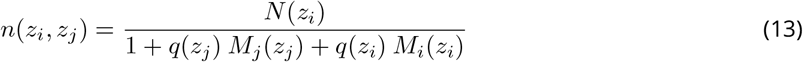

with *i* = 1, *j* = 2 for the growth of phenotype 1 and *i* = 2, *j* = 1 for the growth of phenotype 2. Then, we perform a bifurcation analysis where we determine the ecological equilibria and their stability from the eigenvalues of the Jacobian matrix for a gradient of the total amount of nutrients in the lake *N*_0_ from 0 to 10 mg/m^2^.

Ecological equilibria and evolutionary singularities were computed using Wolfram Mathematica 12.2, and eco-evolutionary simulations were run in R using the lsoda integration method.

## Results

### Ecological dynamics driven by nutrients and light in the absence of evolution

We first report the conditions under which the existence of a macrophyte population is feasible (a positive and stable equilibrium) for a fixed depth (no evolution). As expected, feasibility is possible as long as nutrients and light are sufficient at the specific lake depth considered (cf. Appendix A, eq. A5). More specifically, the strength of light attenuation *w* controls how deep macrophytes survive, since turbidity *W* (*z*) increases with depth *z*. For weak light attenuation, life for the macrophytes is possible even at the bottom of the lake, whereas for strong light attenuation, life for the macrophyte is possible only close to the surface. The other controlling factor, nutrient concentration *N* (*z*), decreases as depth *z* approaches 0, implying that the strength of nutrient diffusion *u* controls how close to the surface macrophytes survive before starving for nutrients. When nutrient diffusion is strong enough, macrophytes survive at the surface, whereas for low nutrient diffusion, macrophytes survive only at the bottom. Very low nutrient diffusion and strong light attenuation preclude life at any depth and lead to macrophyte extinction.

### Evolutionary scenario 1

#### Light and nutrients determine the evolution of a macrophyte’s depth in the lake

In the first scenario, where light is attenuated with depth and nutrients are diffused from the bottom of the lake (Fig.1A), depth position simply determines the quantity of light and nutrients a macrophyte population had access to. As a result, the macrophyte population systematically evolved towards a selected depth (CSS) (Fig. 3A, 4A).

**Figure 3.**
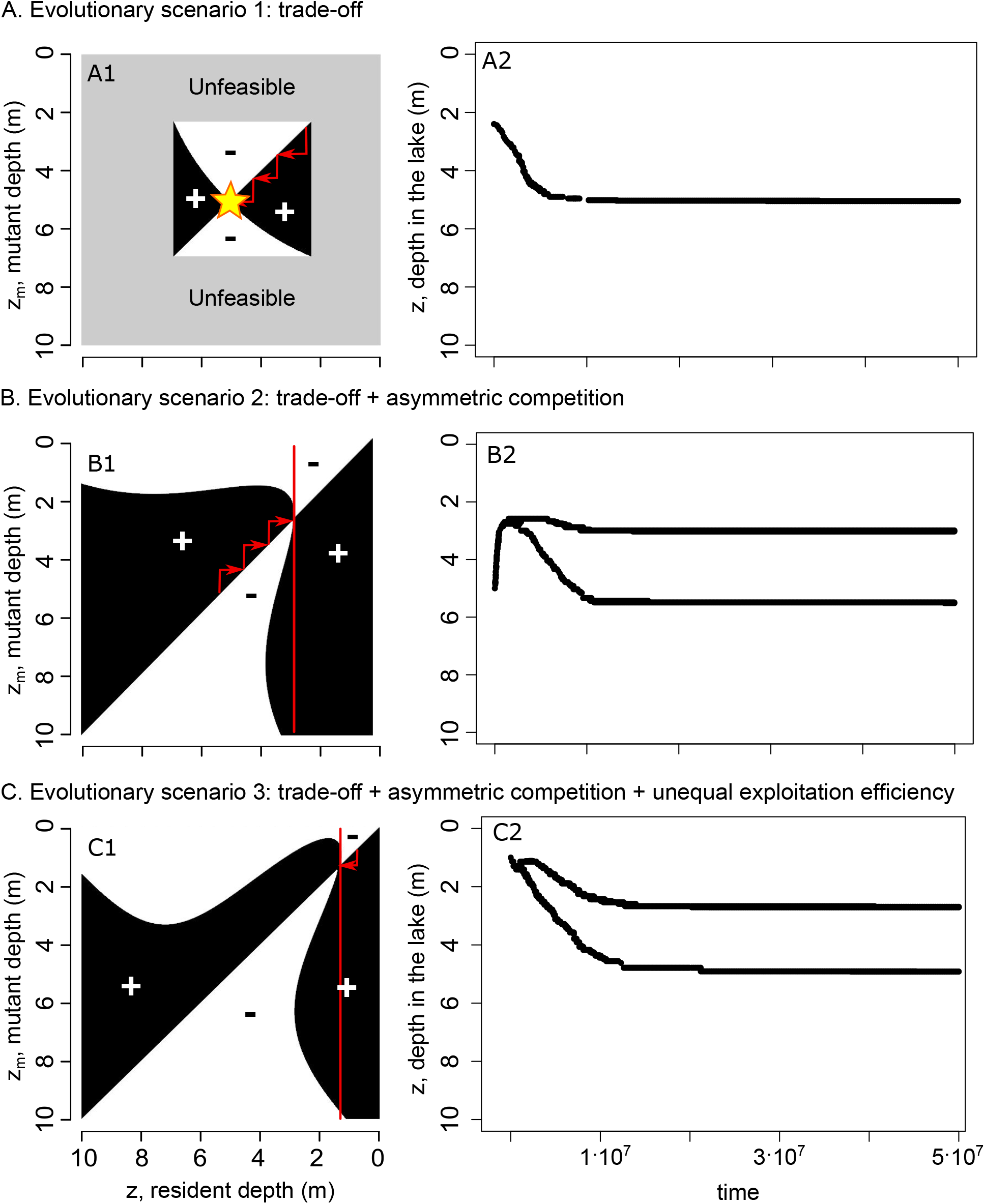
(Left panels): Pairwise Invasibility Plots (PIPs) showing the invasion fitness’ sign depending on resident and mutant traits. Red arrows show the direction of evolution due to selected mutations. Star denotes the Continously Stable Strategy (CSS) Vertical red lines serve as guides to see the positive invasion fitness of mutants around Branching Points (BPs). Grey area denotes the trait values where the population growth is not feasible. (Right panels): Simulation of the eco-evolutionary dynamics corresponding to the red arrows and ES found in the PIPs. A: evolutionary scenario 1 (*u* = 3, *w* = 0.3): the population converges to a selected depth. B: evolutionary scenario 2 (*u* = 5.1, *w* = 0.1): the singularity corresponds to a BP where disruptive selection acts, the population diverges into 2 phenotypes. C: evolutionary scenario 3 (*u* = 4.5, *w* = 0.15): as in scenario 2, diversification occurs.

For example, in our 10 meter deep model lake, with strong light attenuation (*w* = 0.3) and intermediate nutrient diffusion (*u* = 3), no population survives shallower than 2 meters (because of nutrient limitation), and no population survives deeper than 7 meters (because of light limitation, Fig. 3A1). The macrophyte population evolves towards a depth of approximately 5 meters (Fig. 3A). This trait value *z*^*∗*^ is the evolutionary singularity (ES), in other words the point where the selection gradient is null (*s*(*zm, z*) = 0), where the two lines in the PIP cross, where a star is placed (Fig. 3A1). At the ES, no mutant population can invade below or above the resident population. Put differently, in a resident population established between 2 and 5 meters, any mutant population slightly deeper can invade and replace the resident population, whereas in a resident population established between 5 and 7 meters, any mutant population slightly above can invade. Thus, the evolutionary strategy was convergent and stable: it is a selected depth (CSS) towards which the macrophyte population evolves. Here, evolution facilitates the persistence of a population: a population starting at the limit of its feasibility range in the lake evolves further away from the limit. A simulation of the eco-evolutionary dynamics confirmed that the population converged to the ES (Fig. 3A2).

The depth at which the population converges corresponds to the best trade-off between access to light and to nutrients. While the PIP and the associated simulation corresponded to a specific case of nutrient diffusion and light attenuation, Fig. 4A1 shows in a *E*^3^-diagram the effect of nutrient diffusion *u* on the ES. As nutrient diffusion *u* increases, the evolved depth in the lake *z*^*∗*^ decreases. Indeed, as nutrients are more homogeneously distributed, macrophytes evolve closer to the surface where they make the most of the available light.

**Figure 4.**
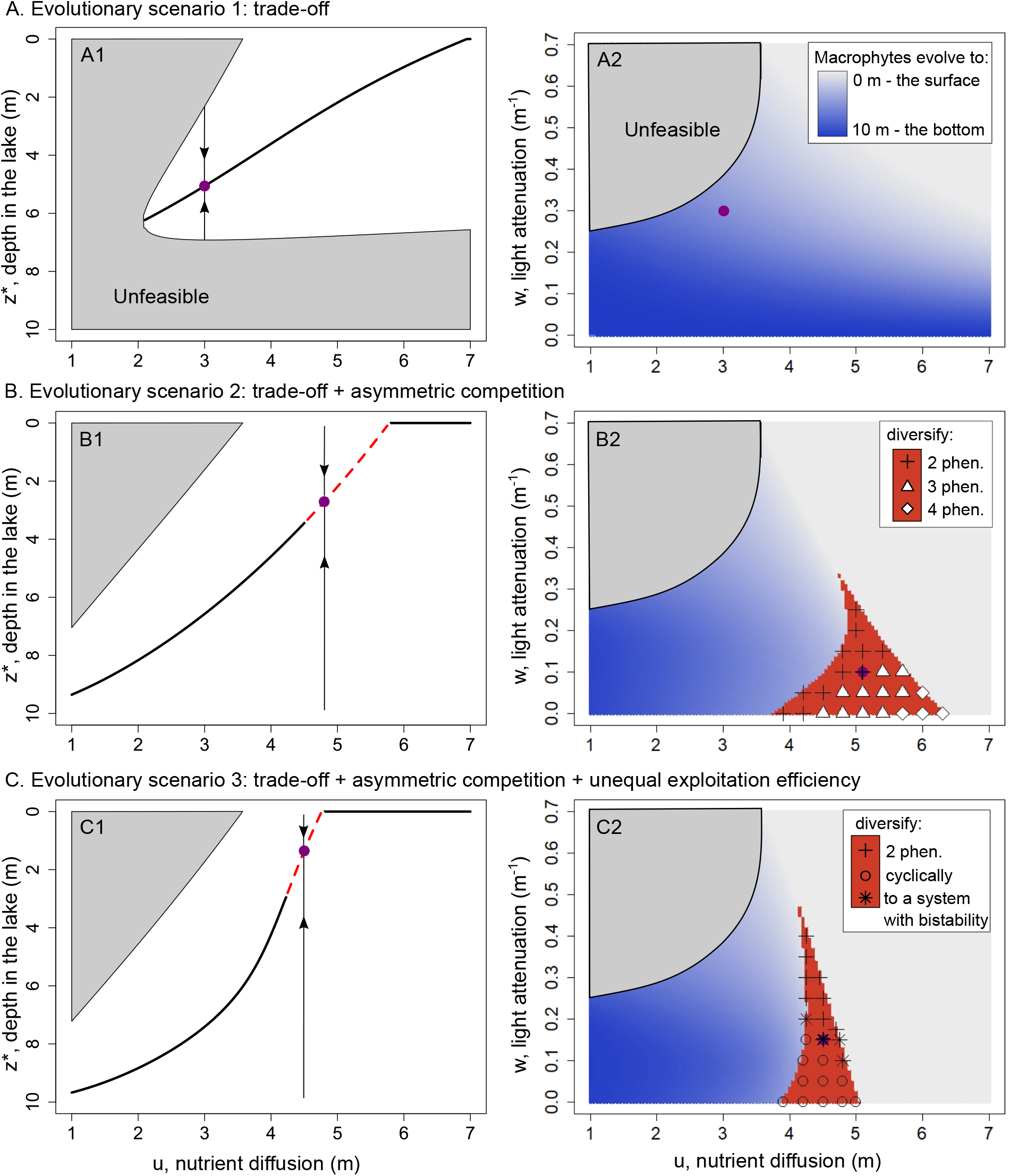
(Left panels) E^3^-diagram showing the ES according to the strength of nutrient diffusion *u*. The population converges to a selected strategy and settles there (CSS, solid black line A1), or converges towards the line then diversifies (BP, dashed red, B1, C1). Arrows show the direction of evolution for phenotypes above or below the ES. (Right panels) Joint effect of nutrient diffusion and light attenuation on the ES. In the blue region of the plot, the population converges towards a selected depth. Diversification (BP) is possible in the red region. Symbols are placed where we ran simulations. Purple dots correspond to simulations of Fig. 3. A: evolutionary scenario 1: The population always converges to a selected depth (CSS). B: evolutionary scenario 2: In most cases, the population converges to a selected depth, but some combinations of nutrient diffusion and light attenuation enable the emergence of polymorphism (B2). C: evolutionary scenario 3: Ecological perturbations after diversification can lead to tipping points (* in C2.)(ES: Evolutionary Singularity, CSS:Continously Stable Strategy, BP: Branching Point)

Finally, we explore the combined effect of nutrient diffusion and light attenuation (Fig. 4A2). This graph is a two dimensional version of the *E*^3^-diagram in Fig. 4A1, where the ES depth *z*^*∗*^ is represented by a shade of blue. For every value of *u* and *w* (Fig. 4A2), the ES is always a selected strategy (CSS). The corners of the figure correspond to the extreme combinations of nutrient and light availability: in the top left corner, low nutrient diffusion and strong light attenuation rendered life impossible. In the bottom left corner, high light availability led the population to evolve to the bottom of the lake (deep blue). In the top right corner, nutrients were strongly diffused so the population could evolve to the surface (light grey). For any intermediate combination, macrophytes evolve to intermediate depths which correspond to the best trade-off between access to light and nutrients. Fig. 4A2 clearly demonstrates that when considering only a light-nutrient trade-off, evolution is not able to lead to an ecosystem with alternative states as the ecosystem is always dominated by a single best-adapted macrophyte population.

### Evolutionary scenario 2

#### Asymmetric competition by shading enables macrophyte diversification

In the second scenario, in addition to the light-nutrient trade-off, we also consider the effect of asymmetry in competition for light. Adding this mechanism enables the macrophyte population to diversify, meaning that more than one phenotype can be supported in the lake.

In Fig. 3B1, the PIP shows how such a diversification arises. The population converges again towards the singularity (positive invasion fitness above the diagonal when the mutant was below the *z*^*∗*^, positive invasion fitness below the diagonal when a mutant was shallower than *z*^*∗*^). However, once at *z*^*∗*^, any mutant invading from a shallower or deeper depth has a positive fitness (the vertical line passing through *z*^*∗*^ is in black regions in the vicinity of *z*^*∗*^). This means that at this point, selection is disruptive. The ES is convergent but invasible and allows the population to diversify (Branching Point, BP). The eco-evolutionary simulation confirm that branching is successful and that two populations with distinct phenotypes evolve from a monomorphic population (Fig. 3B2).

The *E*^3^-diagram Fig. 4B1 however reveals that conditions of diversification occur only for a certain range of nutrient diffusion values (4.5 *< u <* 5.5). Outside of this interval, evolution favors a single phenotype following the pattern found under evolutionary scenario 1: as nutrient diffusion increased, the evolutionary singularity rose closer to the surface.

Fig. 4B2 further shows that diversification is only possible in a subset of light attenuation and nutrient diffusion conditions. Within the branching region, macrophytes diversify into two phenotypes in the most restrictive light and nutrient conditions (crosses situated at the upper left boundary of the branching region), while as conditions became less restrictive the population diversify in more than two phenotypes (towards lower light attenuation and higher nutrient diffusion). For sufficient nutrient diffusion (*u >* 6.2), the population systematically evolves towards the surface, whatever the strength of light attenuation. The effect of the parameter *b* controlling the strength of asymmetry in light competition is presented in detail in Appendix B. The region of diversification changes shape and becomes larger with a higher *b*, and leads to increased diversity at the end of the simulation, clearly emphasizing that asymmetric competition largely drove disruptive selection.

We study whether the diversified ecosystem with multiple phenotypes could show alternative states when responding to a perturbation (Appendix C). In the 25 simulated cases (marked in the diversification subset of Fig. 4B2), whether the system diversified into 2,3 or 4 phenotypes, an increase in nutrient concentration in the lake (*N*_0_) cause only gradual smooth transitions from dominance of a single phenotype to coexistence of multiple phenotypes; it does not exhibit abrupt transitions between alternative stable states (Appendix C Fig. D2).

In conclusion, scenario 2 enables the diversification of the macrophytes, which satisfies the existence of multiple phenotypes that is a necessary condition for the emergence of alternative stable states. However, the combination of the two mechanisms alone is not sufficient for evolution to lead to alternative stable states.

### Evolutionary scenario 3

#### Unequal exploitation efficiencies kick off a positive feedback

In the third scenario, we consider a third additional mechanism, that of a better efficiency in nutrient exploitation when a macrophyte grows close to the bottom of the lake. In such conditions, evolution can lead to a system that diversifies and exhibits alternative stable states.

As in scenario 2, the PIP, eco-evolutionary simulations and *E*^3^-diagram in Fig. 3C1,2 and Fig. 4C1 shows that diversification is possible for a certain range of nutrient diffusion *u*. Similar to Fig. 4B2, Fig. 4C2 shows that diversification occurs in a restricted set of nutrient diffusion and light limitation. However, this set is smaller in scenario 3 compared to scenario 2 and spans different limits. In some cases, after diversification, one of the phenotypes go extinct, and the remaining monomorphic population returns to the convergent singularity where disruptive selection started again, leading to a cyclical diversification behavior (Fig. 5C). In addition, in scenario 3 diversification leads to a maximum of only 2 phenotypes, whereas in scenario 2 we find up to 4 different phenotypes (Appendix C Fig. C1).

**Figure 5.**
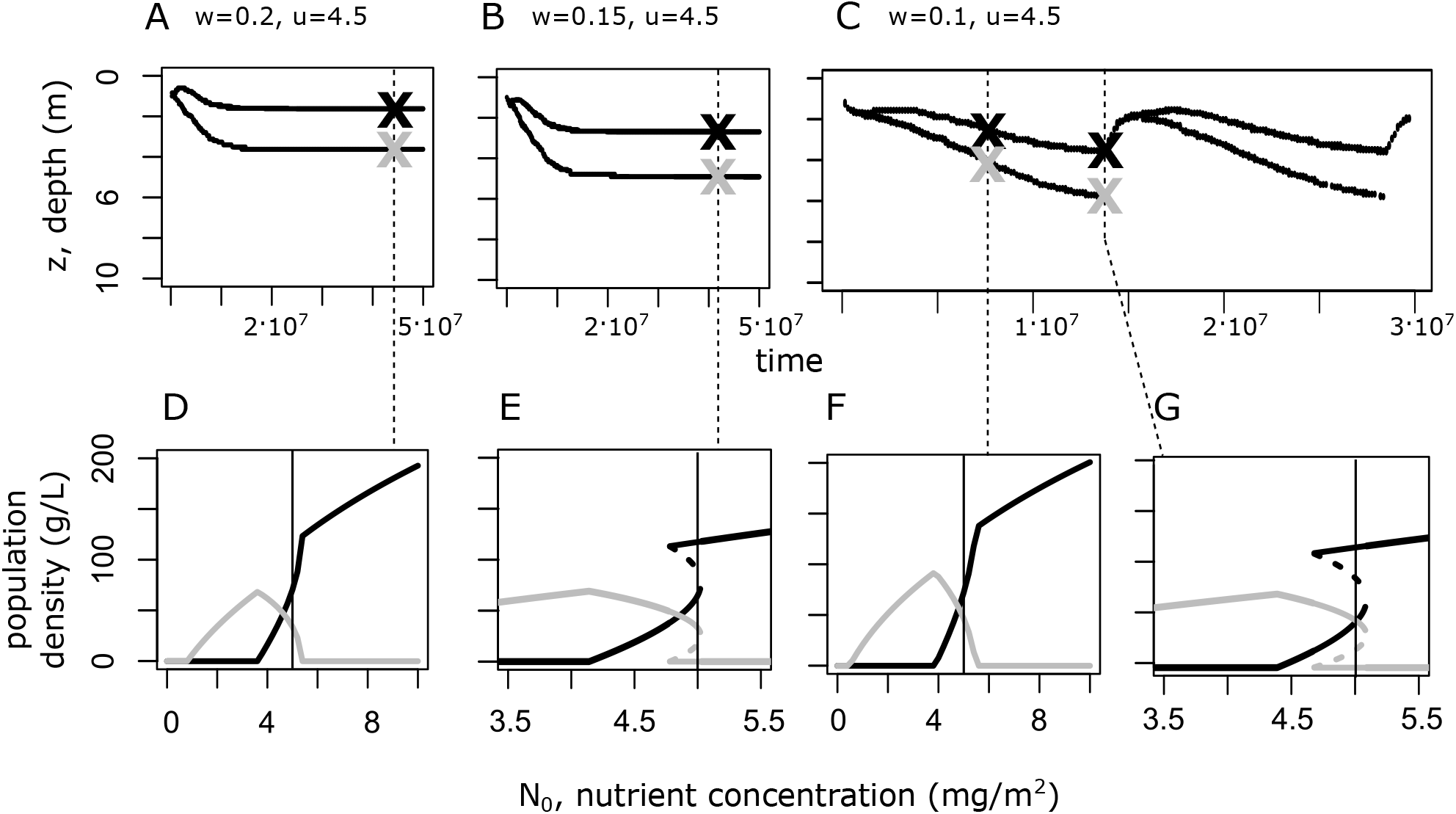
Ecological dynamics with and without alternative states following evolution. A,B,C: Eco-evolutionary simulations of the third evolutionary scenario for different light attenuation conditions. Each line represents the evolution of a phenotype with a specific growth depth *z*. In most cases evolution leads to two dominant phenotypes (A,B), whereas in some cases to cyclical diversification events where after branching one phenotype goes extinct abrupty and diversification repeats itself (C). Crosses mark trait values of *w* and *u* used for the bifurcation analysis of panels D-G. D,E,F,G: Bifurcation analysis of an ecological perturbation of the two evolved macrophytes populations (black and grey lines) from panels A-C. Thin vertical lines recall *N*_0_ at which evolution took place. Solid lines are stable equilibria, dashed lines are unstable equilibria of population density as function of increasing nutrient concentration in the lake *N*_0_. In some cases, the two populations evolve to a depth where they do not respond with tipping points to nutrient enrichment, but with smooth transitions (transcritical bifurcations) from dominance of the deeper macrophyte (gray line) to coexistence and outcompetition by the shallower macrophyte at high nutrient levels (A,D). In other cases, populations evolve at a depth where nutrient enrichment leads to coexistence followed by the deeper macrophyte collapsing through a tipping point (fold bifurcation) when outcompeted by the shallower phenotype. (B,E). In the case of cyclical diversification the response to nutrient enrichment depends on the timing of disturbance: smooth early in the evolutionary cycle (F), or abrupt (through a tipping point) closer to the end of the cycle (G).

In four out of the twelve cases (marked by black stars in Fig. 4C2), we found abrupt transitions between alternative states in response to nutrient enrichment (Fig. 5E). For example, at intermediate nutrient diffusion (*u* = 4.5) and low light attenuation (*w* = 0.15), the populations settled at *z*_1_ *≈* 3 and *z*_2_ *≈* 5 meters (Fig. 5B). For these two phenotypes we find bistability in the range between 4.7 and 5 mg/m^2^ of total nutrient concentration *N*_0_ (Fig. 5E). In the case of cyclic branching (Fig. 5C), we find that the emergence of alternative states depends on the trait values during the evolutionary cycle. During the late stage of the cycle and before the population becomes monomorphic again, alternative states are possible (Fig. 5G), while at the early part of the cycle, no alternative states exist (Fig. 5F). This means that during the evolution of macrophytes the lake can, or not, experience tipping point responses. For the rest of *u* and *w* combinations, we did not find alternative states but smooth transitions (Fig. 5A,D), just like in scenario 2 (Appendix C Fig. C2).

In sum, the third scenario shows a full range of possible eco-evolutionary dynamics, from cases in which a monomorphic population settles at a given depth (CSS), to cases in which disruptive selection can lead to stable or cyclic diversification. In the latter two cases, for a specific range of conditions and depending on the selected depth *z* of the two diversified phenotypes, alternative stable states can emerge.

## Discussion

Tipping point responses between alternative states have been increasingly considered a major risk to ecosystems under global change (Barnosky et al., 2012), but the question of when evolution could lead to their emergence has been largely neglected. In this article, we explored whether the evolution of a macrophyte population in a shallow lake can lead to an ecosystem with tipping points. We used adaptive dynamics to study the evolution of a macrophyte’s vertical position in a lake to find the conditions that permit the emergence of alternative states (Scheffer, Szabo, et al., 2003). We show that, in addition to the light-nutrient trade-off gradient present in a shallow lake, two other mechanisms are necessary. First, asymmetric competition for light (or nutrient, Appendix D) is necessary to induce diversification. Second, unequal exploitation efficiency ensures -but only under some conditions- that evolution can lead to an ecosystem with alternative stable states (Table. 2). In general, our findings corroborate the ecological mechanisms proposed to induce alternative states between floating and submersed macrophytes (Scheffer, Szabo, et al., 2003). Nonetheless, it is unclear whether evolution could have led to alternative states given these mechanisms in the first place. In that sense, our results shed light on how each of these mechanism affects the eco-evolutionary dynamics of macrophytes on their ability to invoke ecosystem tipping point responses.

### Diversification: the necessary condition for the emergence of alternative states

Alternative stable states in a shallow lake are characterized by the dominance of two competing macrophytes that occupy different depths in the water column (ie. submersed and floating macrophytes). Thus, our first objective was to study under which conditions evolution could allow macrophytes to diversify into at least two distinct phenotypes. We hypothesized that the trade-off between access to light and nutrients (ie. parameters *w* and *u* respectively) would determine whether the macrophyte population could diversify. In the case of a weak trade-off (no light attenuation and low nutrient diffusion, or strong light attenuation and strong nutrient diffusion), we expected the population to converge to extreme strategies, that is the surface or the bottom of the lake. By changing the trade-off with the parameters controlling nutrient diffusion and light attenuation (*u* and *w*), we expected that diversification would be possible, as earlier theoretical work showed that in the AD framework, trade-offs between traits can determine the possibility of diversification (Kisdi, 2015; Mazancourt and Dieckmann, 2004). In the case of phytoplankton evolving in a poorly-mixed water column, Klausmeier and Litchman, 2001 and Wickman et al., 2017 found contrasting results (evolution to an optimal depth in the former and diversification in the latter) using different growth functions. In the case of shallow lakes, we expected two types that would grow either at the shallow or deep parts of the lake to resemble the observed floating and submersed macrophytes. However, we found that by only modifying the nutrient diffusion and light attenuation parameters, regardless of the resulting trade-off (*u* and *w*), the macrophyte population never diversifies (Evolutionary scenario 1). Instead, the macrophyte evolves to a single optimal depth, as found in Klausmeier and Litchman, 2001.

The mere existence of a trade-off is thus insufficient to lead to diversification. Instead, we found that asymmetric competition for light through shading of a shallow phenotype to phenotypes growing deeper in the water column enabled diversification (evolutionary scenario 2). This result does not hold only for light as we found similar results also for asymmetric competition for nutrients. This finding resonates with Kisdi, 1999 who showed that the evolutionary dynamics of a continuous trait in a simple asymmetric Lotka-Voltera competition model led to diversification. They further showed that increased asymmetry led to more branching events. In our model we found increased diversity when the strength of asymmetric competition for light (*b*) is high (Appendix B B1). In terrestrial communities, a forest model in which competition for light was asymmetric also led to diversified communities (Falster et al., 2017). Nonetheless, asymmetric competition does not always lead to diversification. We found combinations of nutrient diffusion *u* and light attenuation *w* that lead to a single selected depth defined by the light-nutrient trade-off (mechanism 1). Such convergence to a single depth occurs for strong nutrient diffusion or strong light attenuation (at the right of the diversification region in Fig. 4B2,C2), which corresponds to cases where the incentive to go to the surface overpowers the effects of asymmetric competition. In other cases, convergence to a single depth occurs when environmental conditions are quite limiting, close to the “Unfeasible” area which corresponds to conditions that prevent life at any depth. In these limiting conditions, the biomass of the macrophyte population is not sufficient for the asymmetric competition between the resident and the mutant populations to induce disruptive selection (Doebeli and Dieckmann, 2000; Kisdi, 1999).

While our results show that diversification may lead, in some instances, to cases of alternative stable states, we do not mean that sympatric speciation is needed in any way for these to occur. In particular, note that branching events here could represent other phenomena ranging from the emergence of (within species) polymorphism to the sorting of phenotypes through assembly processes based on pre-existing variability (both intra-specific variability and inter-specific variability). For deeper discussions on how eco-evolutionary dynamics relate to ecological community assembly, see Brännström et al., 2012, Edwards et al., 2018, Leibold et al., in press.

#### Evolution allows for alternative states only for a limited range of conditions

Our main objective was to see under which conditions macrophytes would evolve to having alternative stable states. In the presence of the two mechanisms of a light nutrient trade-off together with asymmetric competition due to shading, evolution leads to diversified phenotypes but not to alternative states (Appendix C Fig. C2). Only in the presence of the third mechanism, that of superior nutrient exploitation efficiency of the deepest phenotype, alternative states emerge and that only for a certain range of nutrient and light availability (evolutionary scenario 3). This results is in line with previous ecological work on alternative stable states. In a water column with a light and nutrient trade-off, and where the nutrient dynamics are explicitly modeled, two phytoplankton populations can exhibit alternative stable states (Huisman and Weissing, 1995). This concurs with Scheffer, Szabo, et al., 2003 where the presence of 3 competitive asymmetries is necessary to incur alternative states: 1) submersed macrophytes have access to more nutrients than floating macrophytes, whereas floating macrophytes have more access to light than submersed macrophytes, 2) floating macrophytes limit the growth of submersed macrophytes by shading them out (light competition *a*(*z*_*m*_, *z*), eq. 10) and 3) submersed macrophytes limit the growth of floating macrophytes by better removing nutrients from the water column (retention efficiency *q*(*z*), eq. 11). Note however that these three asymmetries, though necessary, are not sufficient since only some parameter conditions enable the emergence of alternative stable states (Fig. 4C2).

Although alternative stable states may emerge in our eco-evolutionary model, we do not find that the macrophytes diversify to the two distinct phenotypes, ie. “floating” (*z* = 0) and “submersed” (*z* = *z*_*b*_) typically observed in ponds as modeled by Scheffer, Szabo, et al., 2003. Diversification between light specialists and nutrient specialists has been predicted by theoretical work on the evolution of phytoplankton (Troost et al., 2005), and studies have shown that alternative stable states are possible in models of competition for light and nutrients in aquatic ecosystems (Huisman and Weissing, 1995; Yoshiyama and Nakajima, 2006), under the condition that competitors are well-differentiated in their ability to uptake light and nutrients. In our case, evolved phenotypes are not very different, but only separated by approximately 2 meters in our modeled 10 meter shallow lake (Fig. 3 B2,C2). This relatively small difference between the diversified phenotypes is explained by the trade-off for light and nutrients that allows diversification only within a range of (intermediate) depths. Out of this range, diversification is not possible because shallow phenotypes would lack nutrients, whereas deep phenotypes would lack light. Consequently, we find a specific range of conditions where light and nutrient availability allow the existence of diversified phenotypes (red area in Fig. 4B2,C2) and an even more limited range where the phenotypes constitute alternative states (stars in Fig. 4C2). However, the cyclic branching events offer evidence for the idea that sufficient differentiation is required between the two phenotypes. Indeed, alternative stable states can only occur late in the diversification cycle when the populations are different enough, but not early.

#### Small phenotypic differences can lead to strong positive feedbacks

To some extent it might seem odd that alternative states arise for rather small phenotypic difference between the two phenotypes. Being relatively similar to each other implies that the ability of both phenotypes to acquire nutrients and light is not very different. It is, however, the asymmetric competition due to shading and unequal exploitation efficiencies that are the strongest at only small differences in depth (Fig. 2C,D) that determine the strength of the positive feedback between the two macrophytes and eventually the existence of alternative states. In fact, it appears that the strength of the positive feedback depends not only on the difference in depth (which sets the effect of competition) but also on the relative position of the two macrophytes in the lake (that sets the effect of the light-nutrient opposing gradient). For two phenotypes growing in similar but shallow depths, outcompetition occurs gradually (Fig 4A,D), whereas when the two macrophytes become less similar and grow in deeper depths the resulting asymmetries are strong enough to create positive feedbacks for outcompetition to occur through a tipping point. This is important to note, as our finding contrasts results of other eco-evolutionary models (eg. Kéfi et al., 2007), where typically evolutionary changes of the trait that directly controls the strength of the positive feedback determine the extent of alternative states. In our case, it is the interaction between direct competition and the environmental conditions under which competition acts that determines whether alternative states will emerge.

#### Limitations

Our model is a simplistic representation of macrophyte evolution. Macrophytes have quite complex life cycles (Bakker et al., 2013), reproducing by rhizomes and acquiring nutrients not just by their roots but also by their stems and leaves. Yet, the intermediate depth phenotypes we recovered could resemble pondweeds (*Potamogeton spp*.) and stoneworts (*Charophytes spp*.), which are both submersed macrophytes and have been shown to asymmetrically compete for light and carbon (Van Den Berg et al., 1999). Here, by using the evolving trait, *z*, as the depth at which a macrophyte lives, we simplistically assume that this depth determines the average availability of nutrients and light as well as the average shading and nutrient access of a macrophyte. Clearly, a more realistic formulation would integrate along the whole depth that a macrophyte lives, as floating macrophytes have roots that can grow deep in the water column, and submersed macrophytes absorb nutrients not just from their roots. Alternatively, our evolving depth could also represent diversified phytoplankton layers. Phytoplankton can determine their position on the water column through flagellates or buoyancy and compete strongly for light and nutrients (CS Reynolds, 1984). Our results could therefore complement previous theoretical studies on the evolution of a single or multiple depth phytoplankton layers (Huisman and Weissing, 1995; Klausmeier and Litchman, 2001; Mellard et al., 2011) by showing that two layers of phytoplankton may also represent alternative states. Note that we only considered a lake with a depth of 10m, and changing that depth to 2 or 3 meters might alter the outcome of evolution and for example allow the emergence of floating and submersed macrophytes.

### Conclusion

More broadly, our work can help better understand how evolution works in the context of opposing resource gradients and multiple feedbacks. The opposing gradients of nutrient availability and light limitation that create a trade-off between two resources is not unique to lakes. Nutrient and light trade-offs can be found in seagrass beds (Williams, 1987), or in grasslands selecting for deep roots or higher stems in terrestrial plants (Olff et al., 1990). In wetlands, opposing stress gradients of anoxic to oxygenated soils coexist with a contrasting low to high salinity gradient (Sanderson et al., 2008), and in hilly drylands water availability follows an opposite gradient to erosion stress (Bautista et al., 2007). Studying coexistence in opposing gradients of resources has been done extensively in aquatic systems (Huisman and Weissing, 1995) as well as in terrestrial systems (Falster et al., 2017; HL Reynolds and Pacala, 1993). While evolution is expected to predominantly lead to a single best-adapted phenotype or coexistence, our work shows that considering competitive asymmetries that can be caused by the opposing gradient itself has the potential to allow alternative stable states to emerge by natural selection. Whether such competitive asymmetries are present and could explain the occurrence of alternative stable states in ecosystems around us is a worthy question to ask.

## Acknowledgements

We thank the HPCave center at UPMC-Sorbonne Université (https://hpcave.upmc.fr/) where simulations were performed. Version 3 of this preprint has been peer-reviewed and recommended by Peer Community In Ecology (https://doi.org/10.24072/pci.ecology.100100)

## Data, script and code availability

Scripts used to perform analysis, simulations, and produce figures 3,4,5, and supplementary figures are available online (https://doi.org/10.5281/zenodo.6606311 or https://github.com/ardichvili/Evolutionary_emergence_alternative_stable_states)

## Annexes or Supplementary Information

### Appendix A: Equilibrium stability and feasibility

Solving f(M) = 0 yields three possible equilibria:

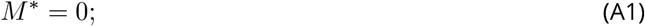

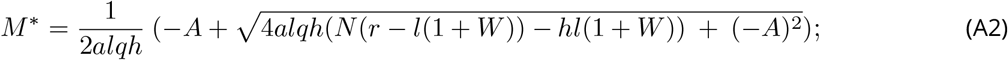

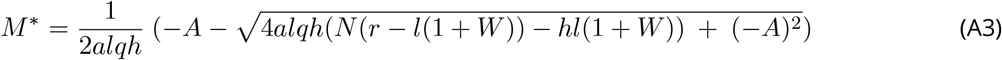

Where

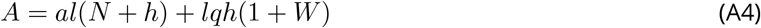

Studying a macrophyte population requires that it exists, ie. that its density is positive. We determined the conditions of feasibility: the combinations of parameters that allow the equilibrium density of macrophytes *M* ^*∗*^ to be positive. We also determined the stability conditions, ie. whether the equilibrium is eventually reached. Local stability requires that *f* ^*′*^(*M* ^*∗*^) *<* 0.

*M* ^*∗*^ in eq. A3 is negative since all parameter values are positive, hence it corresponds to an unfeasible equilibrium.

eq. A2 can be positive if (*N* (*r − l*(1 + *W*)) *− hl*(1 + *W*)) is positive.

This requires

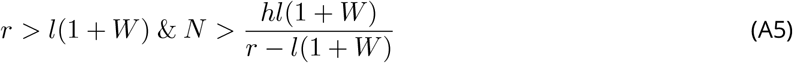

Equation A5 shows that at a given depth *z*, increases in nutrient concentration *N* or in macrophyte growth rate r relaxes the stability conditions. For small nutrient concentrations or negative or low growth rates, the only stable equilibrium is the extinction of the population. Increases in nutrient limitation *h*, mortality *l* or background turbidity *W* could hinder the feasibility and stability of the positive equilibrium.

Since in all three scenarios depth is associated to a trade-off between access to light and nutrient, *N* and *W* vary with depth, *z*.

Since turbidity *W* (*z*) increases with depth *z*, some strength of light attenuation *w* will prevent the growth of macrophytes below a certain depth. In other words, the strength of light attenuation *w* controls how deep macrophytes can survive. For weak light attenuation, life for the macrophytes is feasible at the bottom of the lake. For very strong light attenuation, life for the macrophyte is feasible only close to the surface.

Nutrient concentration *N* (*z*) decreases as depth *z* approaches 0, implying that for some conditions of low nutrient diffusion, macrophytes will not be able to invade above a certain depth. The strength of nutrient diffusion *u* controls how close to the surface a macrophyte can survive before starving for nutrient. If nutrient diffusion is strong enough, the macrophytes can survive at the surface. On the other hand, for a very low diffusion, life is feasible only at the bottom.

Simultaneously low nutrient diffusion and strong light attenuation will preclude life at any depth.

### Appendix B: Effect of the strength of asymmetry in competition for light

In scenario 2 and 3, the introduction of the asymmetric competition for light introduces an additional parameter whose effect can be explored: parameter *b*, which corresponds to the intensity of competition for light. Increased asymmetry in competition for light lead to more diversification events (Fig. B1)

**Figure B1.**
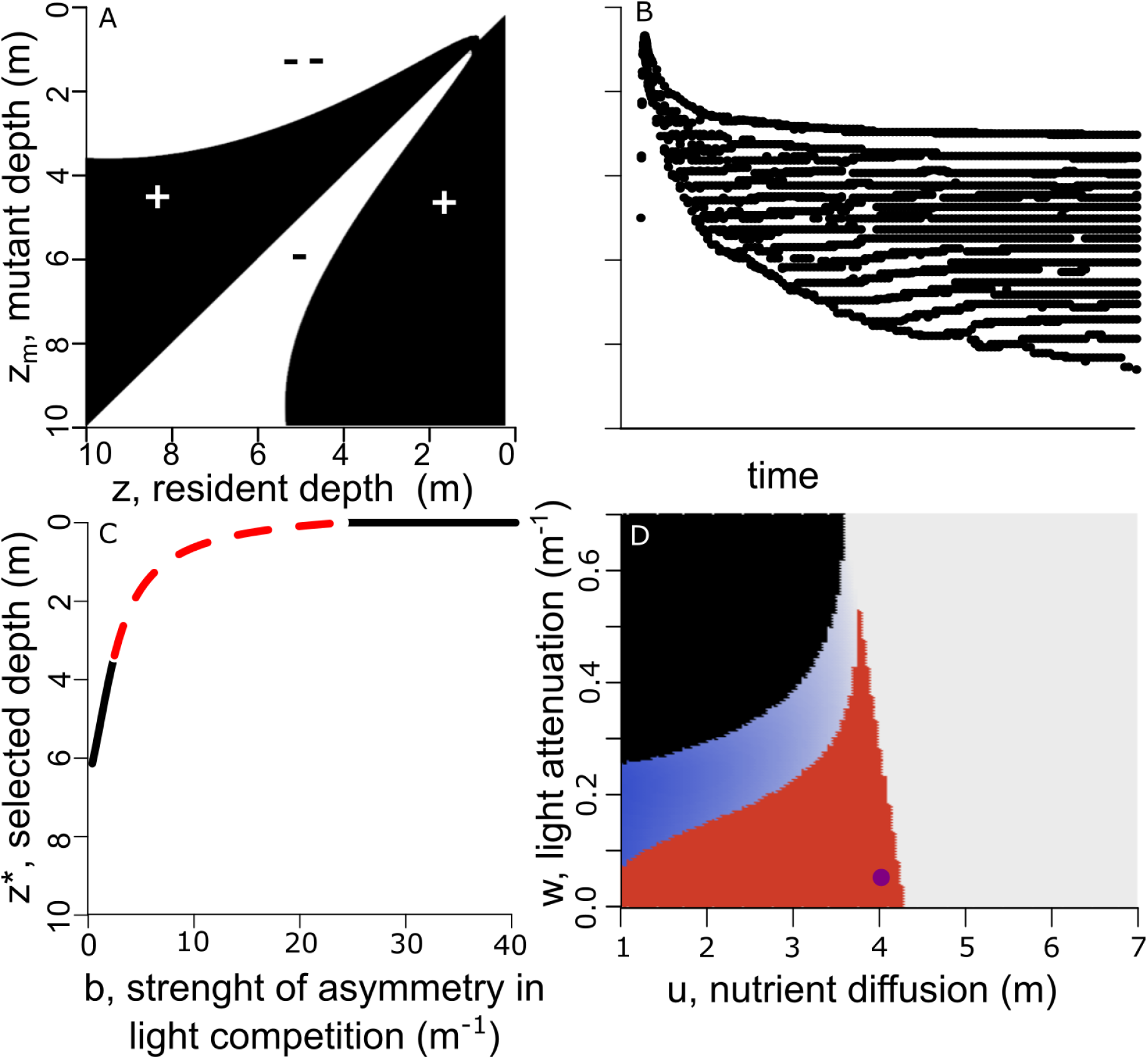
Effect of increased asymmetry in competition for light in Scenario 2. Increased asymmetry can cause more branching events A: PIP for strong nutrient diffusion (*u* = 4), low light attenuation (*w* = 0.05), and strong asymmetry in competition for light (*b* = 5) showing a BP. B: Simulation associated to the conditions in A. More branching events occurred, and coexisting phenotypes live closer together than in Fig.3.B2 C: *E*^3^-diagram for a fixed light attenuation (*w* = 0.2) and fixed nutrient diffusion (*u* = 4). Black solid line: the singularity is a CSS. Red dashed line: the singularity is a BP. As the effect of competition for light increases, the singularity rises closer to the surface. Before it reaches the surface, diversification is possible D: Effect of nutrient diffusion and light attenuation on the singular strategy. Legend is the same as in Fig.4.A2/B2/C2. Relative to the case with weaker asymmetry (Fig.4.B2), the conditions that enable diversification are bigger. However, the population will converge towards the surface more quickly (for *u >* 4.2, whatever the light attenuation)

### Appendix C: Ecological responses of a diversified community in scenario 2

Fig. 4B2. shows that the system can sometimes diversify into more than 2 phenotypes (Fig. C1).

**Figure C1.**
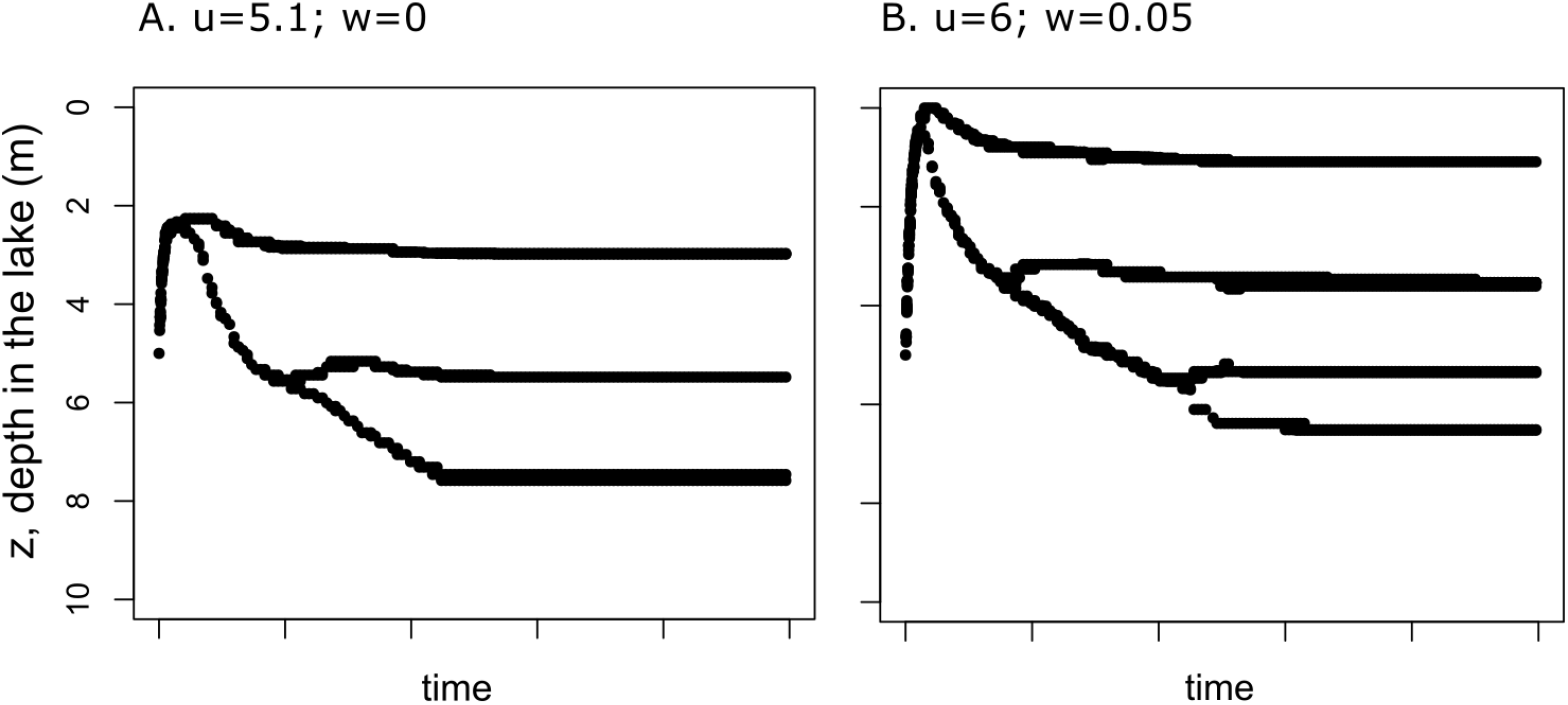
Diversification to more than 2 phenotypes is possible.

Even in diverse communities, theoretical work showed that alternative stable states can exist due to the emergence of clusters of allied species (Nes and Scheffer, 2004). Indeed, in randomly drawn communities in which the interspecific competition coefficients were allowed to vary, some clusters of species in which the intra-cluster competition is lower than the inter-cluster competition formed. The emergence of such “allied” species propagates positive feedback loops and the possibility of alternate stable states. Yet, nutrient enrichment in scenario 2 does not lead to alternate stable states: the transition from the dominance of one phenotype to the other is smooth (Fig. C2).

**Figure C2.**
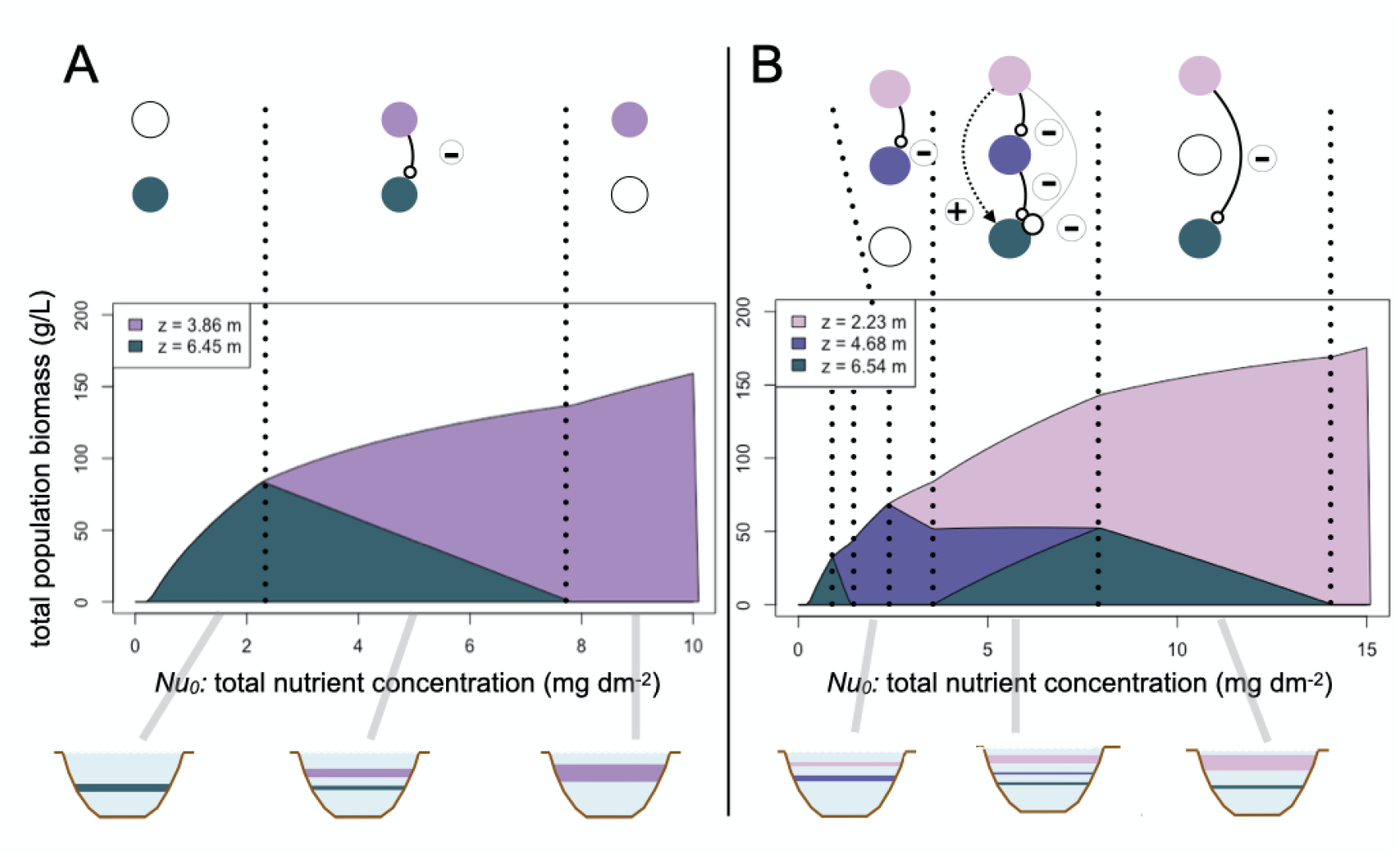
Effect of nutrient enrichment on the abundance of the different present phenotypes in a system that has diversified to two phenotypes (A) and 3 phenotypes (B). Competition networks, cumulated abundances and shallow lake illustrations are shown. No alternate stable states exist. The response to nutrient enrichment is driven by indirect effects (dotted arrows in the competition networks).

### Appendix D: A model with asymmetric competition for nutrient also leads to polymorphism

In a model with asymmetric retention of nutrient, diversification is also possible (Fig. D1)

**Figure D1.**
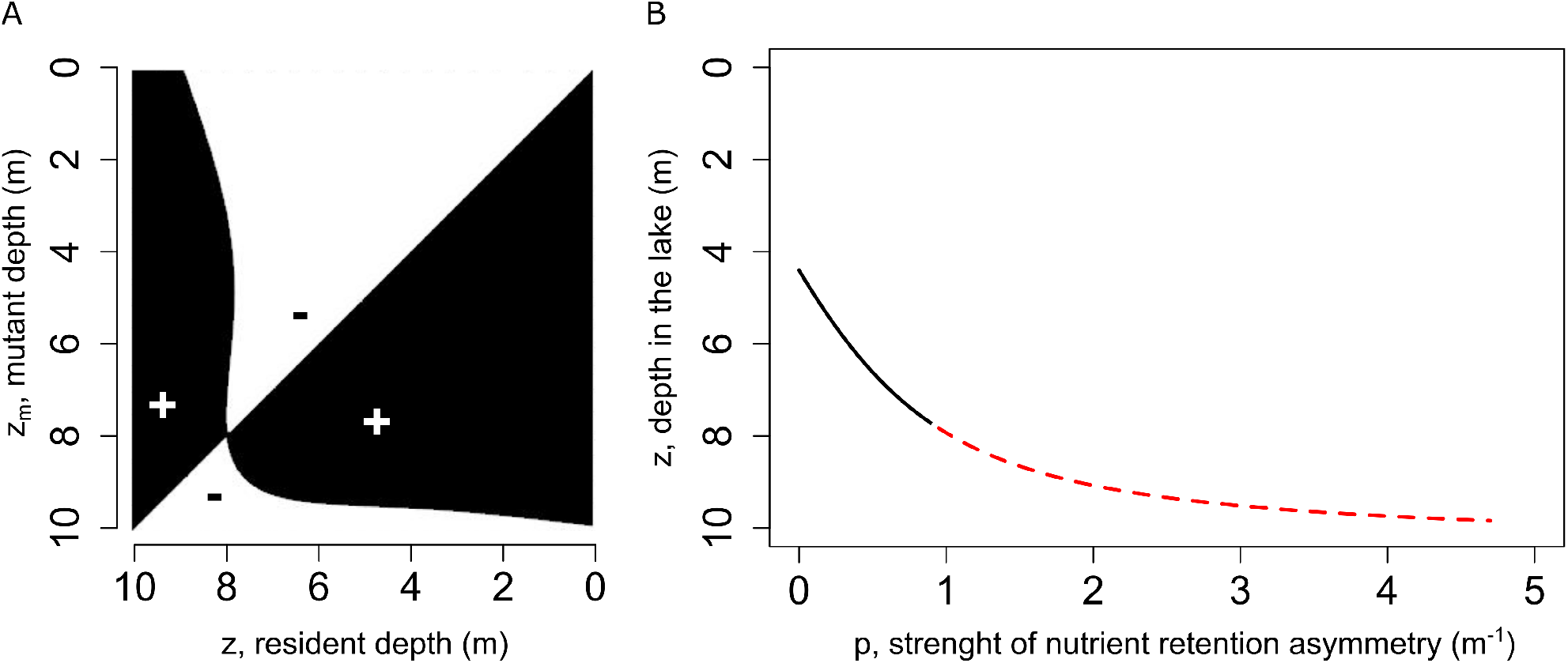
Effect of increased asymmetry in nutrient retention. Increased asymmetry can cause more branching events A: PIP for strong nutrient diffusion (*u* = 5), intermediate light attenuation (*w* = 0.2), and no asymmetry in competition for light (*b* = 0) but asymmetry in competition for nutrient (*p* = 1) showing a BP. B: *E*^3^-diagram for a fixed light attenuation (*w* = 0.2) and fixed nutrient diffusion (*u* = 5). Black solid line: the singularity is a CSS. Red dashed line: the singularity is a BP. As the effect of competition for nutrient increases, the singularity sinks closer to the bottom. Before it reaches the surface, diversification is possible.

## References

Alexander R and J Imberger (Oct. 2008). Spatial distribution of motile phytoplankton in a stratified reservoir: the physical controls on patch formation. Journal of Plankton Research 31, 101–118. ISSN: 0142-7873. https://doi.org/10.1093/plankt/fbn101.

Bakker ES, JM Sarneel, RD Gulati, Z Liu, and E van Donk (2013). Restoring macrophyte diversity in shallow temperate lakes: biotic versus abiotic constraints. Hydrobiologia 710, 23–37. https://doi.org/10.1007/s10750-012-1142-9.

Barnosky AD, EA Hadly, J Bascompte, EL Berlow, JH Brown, M Fortelius, WM Getz, J Harte, A Hastings, PA Marquet, ND Martinez, A Mooers, P Roopnarine, G Vermeij, JW Williams, R Gillespie, J Kitzes, C Marshall, N Matzke, DP Mindell, E Revilla, and AB Smith (June 2012). Approaching a state shift in Earth’s biosphere. en. Nature 486, 52–58. ISSN: 0028-0836, 1476-4687. https://doi.org/10.1038/nature11018.

Bautista S, ÁG Mayor, J Bourakhouadar, and J Bellot (2007). Plant spatial pattern predicts hillslope runoff and erosion in a semiarid mediterranean landscape. Ecosystems 10, 987–998. ISSN: 14329840. https://doi.org/10.1007/s10021-007-9074-3.

Bengfort M and H Malchow (Mar. 2016). Vertical mixing and hysteresis in the competition of buoyant and non-buoyant plankton prey species in a shallow lake. en. Ecological Modelling 323, 51–60. ISSN: 03043800. https://doi.org/10.1016/j.ecolmodel.2015.12.009.

Brännström A, J Johansson, N Loeuille, N Kristensen, TA Troost, RHR Lambers, and U Dieckmann (2012). Modelling the ecology and evolution of communities: a review of past achievements, current efforts, and future promises. en. Evolutionary Ecology Research 14, 601–625.

Chaparro Pedraza PC, B Matthews, L de Meester, and V Dakos (Dec. 2021). Adaptive Evolution Can Both Prevent Ecosystem Collapse and Delay Ecosystem Recovery. en. The American Naturalist 198, E185–E197. ISSN: 0003-0147, 1537-5323. https://doi.org/10.1086/716929.

Chaparro-Pedraza PC and AM de Roos (Mar. 2020). Ecological changes with minor effect initiate evolution to delayed regime shifts. en. Nature Ecology & Evolution 4, 412–418. ISSN: 2397-334X. https://doi.org/10.1038/s41559-020-1110-0.

Cortez MH, S Patel, and SJ Schreiber (Jan. 2020). Destabilizing evolutionary and eco-evolutionary feedbacks drive empirical eco-evolutionary cycles. en. Proceedings of the Royal Society B: Biological Sciences 287, 20192298. ISSN: 0962-8452, 1471-2954. https://doi.org/10.1098/rspb.2019.2298.

Dakos V, B Matthews, AP Hendry, J Levine, N Loeuille, J Norberg, P Nosil, M Scheffer, and L De Meester (Mar. 2019). Ecosystem tipping points in an evolving world. en. Nature Ecology & Evolution 3, 355–362. ISSN: 2397-334X. https://doi.org/10.1038/s41559-019-0797-2.

De Roos AM and L Persson (Oct. 2002). Size-dependent life-history traits promote catastrophic collapses of top predators. en. Proceedings of the National Academy of Sciences 99, 12907–12912. ISSN: 0027-8424, 1091-6490. https://doi.org/10.1073/pnas.192174199.

Dieckmann U and R Law (May 1996). The dynamical theory of coevolution: a derivation from stochastic ecological processes. en. Journal of Mathematical Biology 34, 579–612. ISSN: 0303-6812, 1432-1416. https://doi.org/10.1007/BF02409751.

Doebeli M and U Dieckmann (Oct. 2000). Evolutionary Branching and Sympatric Speciation Caused by Different Types of Ecological Interactions. en. The American Naturalist 156, S77–S101. ISSN: 0003-0147, 1537-5323. https://doi.org/10.1086/303417.

Edwards KF, CT Kremer, ET Miller, MM Osmond, E Litchman, and CA Klausmeier (Dec. 2018). Evolutionarily stable communities: a framework for understanding the role of trait evolution in the maintenance of diversity. en. Ecology Letters 21. Ed. by Becks L, 1853–1868. ISSN: 1461-023X, 1461-0248. https://doi.org/10.1111/ele.13142.

Falster DS, A Brännström, M Westoby, and U Dieckmann (Mar. 2017). Multitrait successional forest dynamics enable diverse competitive coexistence. en. Proceedings of the National Academy of Sciences 114, E2719–E2728. ISSN: 0027-8424, 1091-6490. https://doi.org/10.1073/pnas.1610206114.

Ferriere R and S Legendre (Jan. 2013). Eco-evolutionary feedbacks, adaptive dynamics and evolutionary rescue theory. en. Philosophical Transactions of the Royal Society B: Biological Sciences 368, 20120081. ISSN: 0962-8436, 1471-2970. https://doi.org/10.1098/rstb.2012.0081.

Geritz S, E Kisdi, G Meszéna, and J Metz (Jan. 1998). Evolutionarily singular strategies and the adaptive growth and branching of the evolutionary tree. en. Evolutionary Ecology 12, 35–57. ISSN: 0269-7653, 1573-8477. https://doi.org/10.1023/A:1006554906681.

Gomulkiewicz R and RD Holt (Feb. 1995). When does Evolution by Natural Selection Prevent Extinction? Evolution 49, 201. ISSN: 00143820. https://doi.org/10.2307/2410305.

Gsell AS, U Scharfenberger, D Özkundakci, A Walters, LA Hansson, ABG Janssen, P Nõges, PC Reid, DE Schindler, E Van Donk, V Dakos, and R Adrian (2016). Evaluating early-warning indicators of critical transitions in natural aquatic ecosystems. Proceedings of the National Academy of Sciences 113, E8089–E8095. ISSN: 0027-8424. https://doi.org/10.1073/pnas.1608242113.

Holling CS (Nov. 1973). Resilience and Stability of Ecological Systems. en. Annual Review of Ecology and Systematics 4, 1–23. ISSN: 0066-4162. https://doi.org/10.1146/annurev.es.04.110173.000245.

Huisman J and FJ Weissing (Oct. 1995). Competition for Nutrients and Light in a Mixed Water Column: A Theoretical Analysis. en. The American Naturalist 146, 536–564. ISSN: 0003-0147, 1537-5323. https://doi.org/10.1086/285814.

Intergovernmental Science-Policy Platform on Biodiversity and Ecosystem Services I (Nov. 2019). Summary for policymakers of the global assessment report on biodiversity and ecosystem services. en. Tech. rep. IPBES secretariat, Bonn, Germany. https://doi.org/10.5281/ZENODO.3553579.

Kéfi S, M Rietkerk, M van Baalen, and M Loreau (May 2007). Local facilitation, bistability and transitions in arid ecosystems. en. Theoretical Population Biology 71, 367–379. ISSN: 00405809. https://doi.org/10.1016/j.tpb.2006.09.003.

Kisdi E (Mar. 1999). Evolutionary Branching under Asymmetric Competition. en. Journal of Theoretical Biology 197, 149–162. ISSN: 00225193. https://doi.org/10.1006/jtbi.1998.0864.

Kisdi E (Apr. 2015). Construction of multiple trade-offs to obtain arbitrary singularities of adaptive dynamics. en. Journal of Mathematical Biology 70, 1093–1117. ISSN: 0303-6812, 1432-1416. https://doi.org/10.1007/s00285-014-0788-5.

Klausmeier CA and E Litchman (Nov. 2001). Algal games: The vertical distribution of phytoplankton in poorly mixed water columns. en. Limnology and Oceanography 46, 1998–2007. ISSN: 00243590. https://doi.org/10.4319/lo.2001.46.8.1998.

Knowlton N (Dec. 1992). Thresholds and Multiple Stable States in Coral Reef Community Dynamics. en. American Zoologist 32, 674–682. ISSN: 0003-1569. https://doi.org/10.1093/icb/32.6.674.

Kondoh M (Feb. 2003). Foraging Adaptation and the Relationship Between Food-Web Complexity and Stability. Science 299, 1388–1391. ISSN: 00368075, 10959203. https://doi.org/10.1126/science.1079154.

Leibold MA, L Govaert, N Loeuille, L De Meester, and MC Urban (in press). Evolution and community assembly across spatial scales. Annual Review of Ecology, Evolution, and Systematics.

Loeuille N (Dec. 2010). Influence of evolution on the stability of ecological communities: Evolution and stability of communities. en. Ecology Letters 13, 1536–1545. ISSN: 1461023X. https://doi.org/10.1111/j.1461-0248.2010.01545.x.

Mazancourt C de and U Dieckmann (Dec. 2004). Trade-Off Geometries and Frequency-Dependent Selection. en. The American Naturalist 164, 765–778. ISSN: 0003-0147, 1537-5323. https://doi.org/10.1086/424762.

Mellard JP, K Yoshiyama, E Litchman, and CA Klausmeier (2011). The vertical distribution of phytoplankton in stratified water columns. Journal of Theoretical Biology 269, 16–30. ISSN: 00225193. https://doi.org/10.1016/j.jtbi.2010.09.041.

Metz J, R Nisbet, and S Geritz (June 1992). How should we define ‘fitness’ for general ecological scenarios? en. Trends in Ecology & Evolution 7, 198–202. ISSN: 01695347. https://doi.org/10.1016/0169-5347(92)90073-K.

Mumby PJ, A Hastings, and HJ Edwards (Nov. 2007). Thresholds and the resilience of Caribbean coral reefs. en. Nature 450, 98–101. ISSN: 0028-0836, 1476-4687. https://doi.org/10.1038/nature06252.

Nes EH van, BM Arani, A Staal, B van der Bolt, BM Flores, S Bathiany, and M Scheffer (Dec. 2016). What Do You Mean, ‘Tipping Point’? en. Trends in Ecology & Evolution 31, 902–904. ISSN: 01695347. https://doi.org/10.1016/j.tree.2016.09.011.

Nes EH van and M Scheffer (Aug. 2004). Large Species Shifts Triggered by Small Forces. en. The American Naturalist 164, 255–266. ISSN: 0003-0147, 1537-5323. https://doi.org/10.1086/422204.

Olff H, JV Andel, and JP Bakker (1990). Biomass and Shoot/Root Allocation of Five Species from a Grassland Succession Series at Different Combinations of Light and Nutrient Supply. Functional Ecology 4, 193. ISSN: 02698463. https://doi.org/10.2307/2389338.

Reynolds CS (1984). The ecology of freshwater phytoplankton. eng. Reprinted. Cambridge Studies in ecology. Cambridge: Univ. Press. ISBN: 978-0-521-28222-2 978-0-521-23782-6.

Reynolds HL and SW Pacala (Jan. 1993). An Analytical Treatment of Root-to-Shoot Ratio and Plant Competition for Soil Nutrient and Light. en. The American Naturalist 141, 51–70. ISSN: 0003-0147, 1537-5323. https://doi.org/10.1086/285460.

Sanderson JS, NB Kotliar, and DA Steingraeber (2008). Opposing environmental gradients govern vegetation zonation in an intermountain playa. Wetlands 28, 1060–1070. ISSN: 02775212. https://doi.org/10.1672/07-111.1.

Scheffer M, S Szabo, A Gragnani, EH van Nes, S Rinaldi, N Kautsky, J Norberg, RMM Roijackers, and RJM Franken (Apr. 2003). Floating plant dominance as a stable state. en. Proceedings of the National Academy of Sciences 100, 4040–4045. ISSN: 0027-8424, 1091-6490. https://doi.org/10.1073/pnas.0737918100.

Scheffer M (1998). Ecology of Shallow Lakes. English. OCLC: 853265766. Dordrecht: Springer Netherlands. ISBN: 978-1-4020-3154-0.

Scheffer M, S Carpenter, JA Foley, C Folke, and B Walker (Oct. 2001). Catastrophic shifts in ecosystems. en. Nature 413, 591–596. ISSN: 0028-0836, 1476-4687. https://doi.org/10.1038/35098000.

Scheffer M, R Vergnon, JHC Cornelissen, S Hantson, M Holmgren, EH van Nes, and C Xu (Aug. 2014). Why trees and shrubs but rarely trubs? en. Trends in Ecology & Evolution 29, 433–434. ISSN: 01695347. https://doi.org/10.1016/j.tree.2014.06.001.

Staver AC, S Archibald, and SA Levin (Oct. 2011). The Global Extent and Determinants of Savanna and Forest as Alternative Biome States. en. Science 334, 230–232. ISSN: 0036-8075, 1095-9203. https://doi.org/10.1126/science.1210465.

Troost TA, BW Kooi, and SALM Kooijman (Sept. 2005). Ecological Specialization of Mixotrophic Plankton in a Mixed Water Column. en. The American Naturalist 166, E45–E61. ISSN: 0003-0147, 1537-5323. https://doi.org/10.1086/432038.

Van Den Berg MS, M Scheffer, E Van Nes, and H Coops (1999). Dynamics and stability of Chara sp. and Potamogeton pectinatus in a shallow lake changing in eutrophication level. Hydrobiologia 408-409, 335–342. ISSN: 00188158. https://doi.org/10.1007/978-94-017-2986-4_37.

Wickman J, S Diehl, B Blasius, CA Klausmeier, AB Ryabov, and Å Brännström (Apr. 2017). Determining Selection across Heterogeneous Landscapes: A Perturbation-Based Method and Its Application to Modeling Evolution in Space. en. The American Naturalist 189, 381–395. ISSN: 0003-0147, 1537-5323. https://doi.org/10.1086/690908.

Williams SL (1987). Competition between the seagrasses Thalassia testudinum and Syringodium filiforme in a Caribbean lagoon. Marine Ecology Progress Series 35, 91–98. ISSN: 01718630. https://doi.org/10.3354/meps035091.

Yoshiyama K and H Nakajima (Feb. 2006). Catastrophic shifts in vertical distributions of phytoplankton The existence of a bifurcation set. en. Journal of Mathematical Biology 52, 235–276. ISSN: 0303-6812, 1432-1416. https://doi.org/10.1007/s00285-005-0349-z.

